# Development and Analysis of Multiscale Models for Tuberculosis: From Molecules to Populations

**DOI:** 10.1101/2023.11.13.566861

**Authors:** Pariksheet Nanda, Maral Budak, Christian T. Michael, Kathryn Krupinsky, Denise E. Kirschner

## Abstract

Although infectious disease dynamics are often analyzed at the macro-scale, increasing numbers of drug-resistant infections highlight the importance of within-host modeling that simultaneously solves across multiple scales to effectively respond to epidemics. We review multiscale modeling approaches for complex, interconnected biological systems and discuss critical steps involved in building, analyzing, and applying such models within the discipline of model credibility. We also present our two tools: CaliPro, for calibrating multiscale models (MSMs) to datasets, and tunable resolution, for fine- and coarse-graining sub-models while retaining insights. We include as an example our work simulating infection with *Mycobacterium tuberculosis* to demonstrate modeling choices and how predictions are made to generate new insights and test interventions. We discuss some of the current challenges of incorporating novel datasets, rigorously training computational biologists, and increasing the reach of MSMs. We also offer several promising future research directions of incorporating within-host dynamics into applications ranging from combinatorial treatment to epidemic response.

## 1 Introduction

Our collective understanding of biology has become more complex with the advent of tools that allow for dissection and deeper study. In addition, the interconnectedness of components has led to new fields of study such as complex systems and systems biology. Computational and mathematical models have attempted to follow suit with the creation of models that have grown in detail, mechanisms, and complexity.

Multiscale models (MSMs) that capture biological processes occurring over multiple physiological and time scales are an approach that has evolved in concert over the past two decades to help address open problems in many areas of health such as cancer and infectious diseases [21, 34, 38, 46, 85, 113]. With the advent of MSMs has come the necessity to find creative ways to build, link, solve, analyze, and apply them. The ultimate goal of which is to provide useful predictions to biologists and clinicians who can then take the next steps to uncover more details about the systems under study. In an ideal fashion, this is an iterative process with well-established collaborations between wet-lab and theoretical scientists.

Our lab has focused on exactly these tasks for MSMs related to uncovering details for infection with *Mycobacterium tuberculosis* (Mtb). Several questions and features naturally arise as part of these processes. We must first decide at which scale we should start building MSMs. Our approach has been to start at the middle / mesoscale and build out. It is not always obvious which modeling framework is best suited to capture representative datasets, and multiple formulations may be necessary before settling on which is best for addressing open questions (e.g. agent-based models or deterministic models such as ODEs) [36]. We consider the mesoscale as the scale at which most data are available and also the scale that is driving key biological questions. Once a model is built, both model calibration and validation steps are necessary to ensure that model outputs reflect the biological system under study. As part of our discovery process, we also created tools for performing calibration and validation of MSMs for complex biological systems [52, 99] against different modes of spatial and temporal datasets.

After the meso-scale model has been developed, decisions need to be made about which specific physiologic scale(s) to consider next, and which modeling framework to use. A key choice is: should the MSM be developed “north–south” (i.e. higher or lower physiological scales of interest) or developed “east–west” (i.e. adding more physiological compartments occurring at the same scale). Fig. 1 indicates physiological expansions reflecting this choices of MSM “directionality”. The questions of interest usually drive these decisions.

**Fig. 1.**
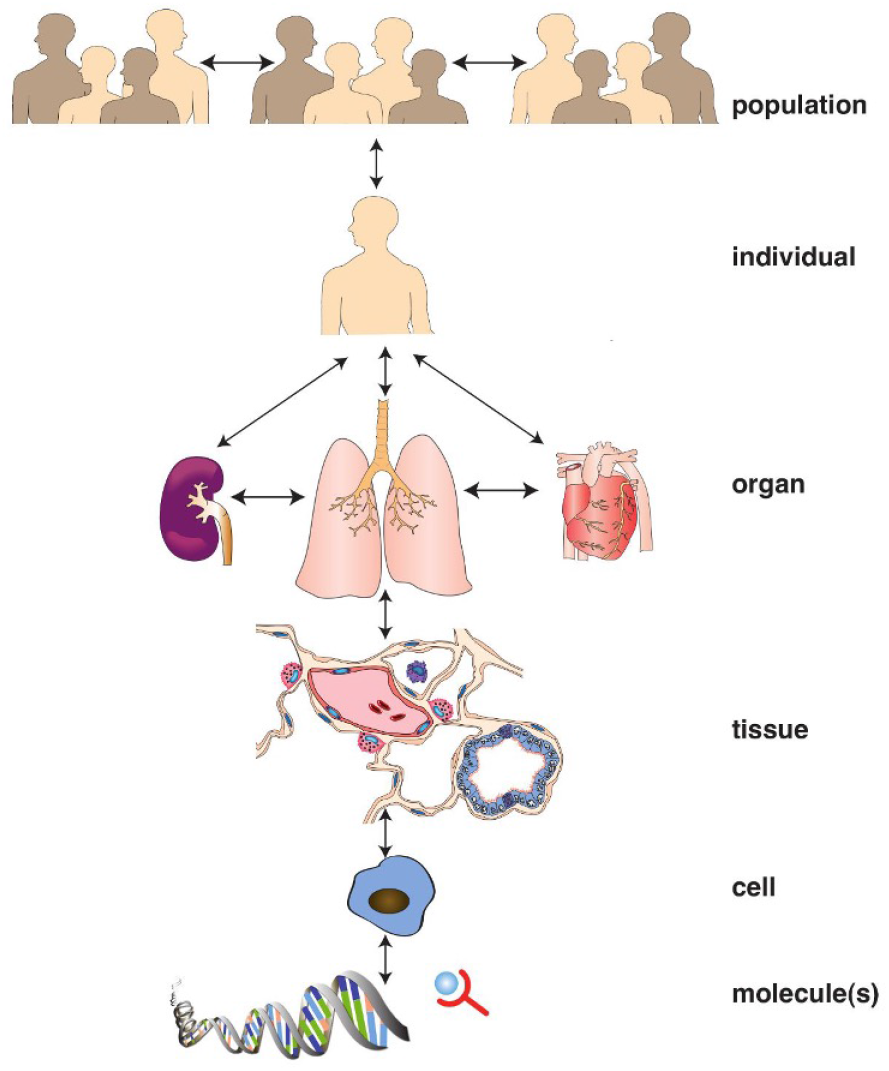
Multiple physiological scales in human health. This also indicates an example of north–south and east– west model expansion from a mesoscale, which, here, is the lungs.

When new scales are added, a required step, that is not yet standardized, is the linking of model scales [61]. Linking independent model components that operate at different temporal or spatial scales, specifically when model components use different model formulations, requires explicit consideration. Some simple ways of linking include: passing parameters across scales, creating lookup tables that send information across scales, and allowing variables within compartmental models to cross to other compartments with appropriate volumetric scaling. Several groups have developed frameworks to modularize models to improve model linking [12, 48, 49, 58, 62, 71, 123]. Additionally, the multiscale modeling platforms CompuCell3D [119], Simmune [81, 114], PhysiCell [40], and Virtual Cell [18, 109] all provide tools for model linking.

Once MSMs are created, analyses of models are necessary to calibrate, validate and also study dynamics to make useful predictions. When MSMs were initially being developed, there were no tools available to study these highly complex and usually-hybrid models. Over the past 20 years, our group has developed and honed a number of approaches for the analysis and study of MSMs. When building models, a key step is the parameterization and calibration of models to available datasets for the biological system under study. Recently, we have developed a novel approach to calibrate complex models to biological datasets [52, 86]. Briefly, a model parameter space is adjusted to acceptable ranges based on pass and fail criteria provided by a user and based on available knowledge of, and specific datasets for, the corresponding biological system. Whether the model is deterministic or stochastic, our framework, called CaliPro, provides a step-by-step process towards this goal. This approach fits into a larger framework of approximate Bayesian computing (ABC). We highlight the strengths and weaknesses of both approaches herein and in previous work [86]. After models are calibrated, MSMs can be analyzed to determine what mechanisms are driving model outputs [73]. Our group has used an approach for performing sensitivity analysis that can identify parameters that represent biological mechanisms within the system under study. This approach, in some sense, can identify bifurcation-like parameters that distinguish system dynamics and steady state behaviors. Once these key parameters are identified, they can be used in a number of ways for MSM development as well as to assist with reducing model complexity, which is necessary as MSMs grow to computationally intensive sizes. We refer to this idea of model reduction that preserves the biology of the system, rather than the mathematics, while simultaneously reducing computational complexity and speed, as tunable resolution [61]. We walk through an example of how this approach works herein.

Over the past 20 years, many MSM researchers have participated as part of a consortium to create a set of *10 Rules* for model credibility that are best practices for model development and testing, derived from engineering principles [26]. The 10 rules are as follows: 1: Define context clearly; 2: Use appropriate data; 3: Evaluate within context; 4: List limitations explicitly; 5: Use version control; 6: Document adequately; 7: Disseminate broadly; 8: Get independent reviews; 9: Test competing implementations; 10: Conform to standards. These rules have broad interpretability and may be model context dependent, so we highlight how we have addressed them. Finally, defining model outputs is a key step that allows our model to both match datasets for the biological system, and also from targeted predictions about mechanisms that are not currently observable in the wet lab. Consequently, well-designed outputs can lend insights into the next best experiments to perform or can direct investigations into new avenues. Similarly, implementing therapies, such as vaccines or other interventions within MSMs, either by inclusion of a new scale in the model or by integration directly into the existing MSM framework, can provide new routes to investigate therapies. Virtual clinical trails can aid in narrowing the design space for drug regimens, optimizing vaccine routes, etc.. These interventions are typically rolled out and tested at a single biological scale, but using a MSM approach, we can simultaneously observe effects at multiple scales; while some interventions seem to be successful at, for example, a tissue scale, when examined with a MSM at an entire population those results may, or importantly, may not hold true. This nicely follows the expert ideas of scalability as outlined by Professor Simon Levin in his MacArthur award paper [64].

Within this review, we explain in detail how we have addressed each of these approaches with the development of MSMs using our work in the area of *M. tuberculosis* infection. We provide detailed examples of each of the points raised above as we walk through the development of MSMs that range from molecular up to the population scales. Along the way, we offer insights into how we made choices and applied approaches to glean the most from each model and what open questions remain to be addressed.

## 2 Building Multiscale Models

We have developed several models of Tuberculosis (TB) disease progression and outline here our guiding decisions behind model formulations, linking scales, and adding scales to previously introduced models, model parameter sampling, and model calibration.

TB infectious disease is caused by infection with the bacterium *Mycobacterium tuberculosis* (Mtb). In humans, TB is characterized by a large latently infected population, highlighting the need to identify immune response drivers of outcomes, including Mtb elimination, latent control, and infectious, clinically active disease.

The immune response to Mtb inhalation in the lungs is characterized by the formation of granulomas — lesions comprised of immune cells and bacteria — that physically contain and immunologically restrain infection (Fig. 2). A characteristic feature of most granulomas is a collection of dead cells and debris that is referred to as a necrotic core or caseum, which protects non-replicating Mtb trapped within, allows long-term persistence, and is also an obstacle to treatment. The challenge of modeling Mtb infection is the involvement of multiple tissues and organs, and the paucity of biologically relevant animal models that replicate the spectrum of disease and granuloma physiology found in humans [10]; in particular non-human primate (NHP) animal models are necessary [10]. Thus, multiscale mathematical and computational models play an important role by representing the spatial, temporal, compartmental, and multi-resolution aspects of global and local dynamics of TB disease [36].

**Fig. 2.**
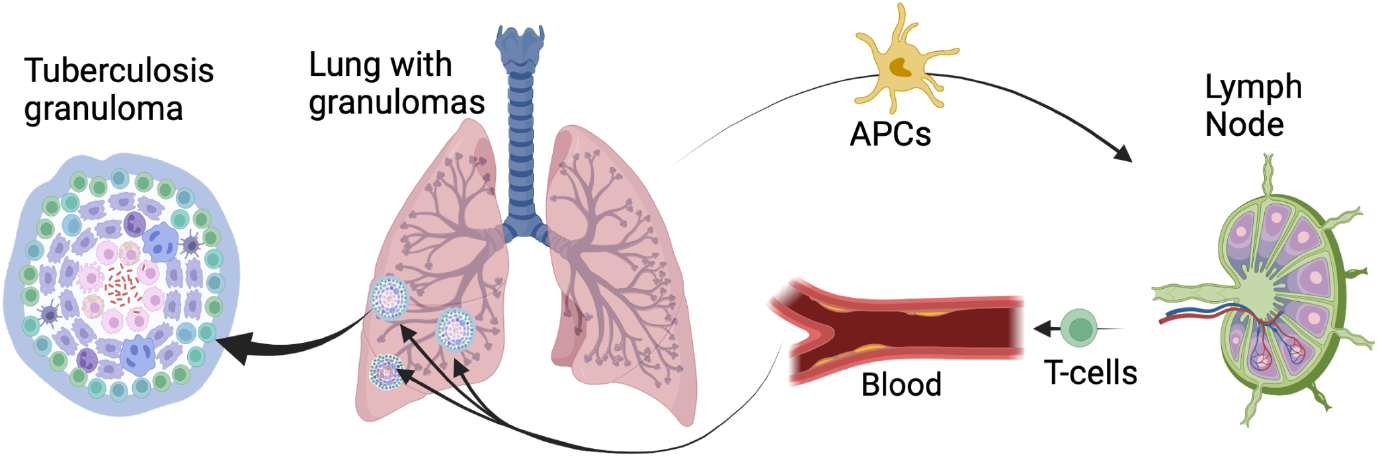
Biological schematic of pulmonary tuberculosis disease. Infection begins when Mtb (red bacterial rods) are inhaled into lungs and phagocytosed by macrophages (purple cells) where Mtb then replicates inside of, and infects, macrophages (pink cells). The immune system creates a physical barrier to infection by forming granulomas around the infection loci comprising macrophages and an outer T-cell cuff (green cells). Infection involves several organs and the T cell adaptive immune response is triggered within lymph nodes by antigen presenting cells (APCs) that migrate from the lungs. Primed T cells migrate through blood to the lung granulomas. In cases of higher infection severity, Mtb disseminates and forms new granulomas.

### 2.1 Formulating Models of Tuberculosis and Linking Scales

Our first TB model used the framework of mathematical modeling with nonlinear ordinary differential equations (ODEs) [126]. This type of model was used to match the non-spatial nature of available experimental datasets: bronchoalveolar lavage fluid containing cellular components, cytokines, and whole, ground lungs of mice [126]. To extend the model beyond the lung organ, we incorporated a lung draining lymph node that primes and activates the adaptive immune response. It was formulated using an additional compartmental ODE model [76, 77]. Although this compartmental model captured cellular migration between the infected lung and lymph node, it still lacked a spatially-related response delay between the site of infection and generation of adaptive immunity. Using only temporal data, we created a spatiotemporal model using a metapopulation model consisting of a coarse grid of compartments within the lung that solved sets of ODEs of cell populations [37]. This model used simpler ODEs within each grid compartment and defined new terms for migration across grid compartments. Next, in a step toward capturing both the states of active disease progression and immune control with granulomas, we represented the macrophage innate immune response using partial differential equations (PDEs) of an Mtb chemokine to capture the growing boundary of a granuloma [35]. Although this coarse-grained spatial model was able to capture disease states of Mtb clearance and initial stages of control, it did not produce latency due to missing adaptive immune cells that are present within granulomas. Finally, focusing on cells as agents, we created an agent-based model (ABM) *GranSim* that captures granuloma formation, damaged necrotic tissue, stochastic cellular behaviors, and molecular diffusion [110]. Spatial agent-based models are defined typically as hybrid models that capture both discretized ODEs / PDEs and other computational formulations usually over both fast and slow timescales. A grid where simulations occur is also defined as well as rules occurring between agents, which in the case of *GranSim* are immune cells, dead tissue, and bacteria. We chose this ABM representation to address questions arising from limitations in prior models and to capture higher-order effects. Importantly, we did a study where we compared the different approaches for modeling and performed analyses comparing what we learned to identify strengths and weaknesses of each approach [36].

Most complex biological systems are difficult to describe using a single spatiotemporal scale: population dynamics and demographics are influenced by individual organisms, whole-individual outcomes are impacted by functionality of organs, organs by tissues, tissues by cells, and so on. Conversely, individual behaviors will be pressured by population dynamics; one tissue may be affected by the functionality of an organ elsewhere in the body, a cell may respond to signals present in a tissue, etc. Consequently, we must ensure to appropriately exchange information between scales to extend the applicable context of our model. We call this process *linking scales*. Because larger scales are characterized by interactions of smaller-scale components, the process of distinguishing scales is somewhat subjective. Hallmarks of larger scales include (i) being aggregates of many similar constituent pieces and (ii) having scientifically-relevant measurable features or emergent behaviors. Many frameworks exist to identify, define, and couple known individual-scale components [12, 48, 49, 58, 62, 71, 123], but defining the distinction between scales is often a system-specific task, with different relevant features defining a larger-scale system. When separating time scales, a common hallmark used to delineate fast from slow is that evolution of a slow timescale is often the drift of a quasi steady-state approximation of the fast timescale [24, 102]. Linking scales exchanges information between them to match measurable biological features or emergent behaviors and yet must sync them computationally and/or mathematically.

Choosing to represent an additional scale should balance biological detail against additional computational burden. When modeling granulomas, we have had success representing mechanisms at a particular scale once we could match biological experiments operating at that scale [11, 15, 28, 30, 42, 66, 72, 75, 78, 94, 95, 118, 127]. Our ability to directly include multi-scale and multi-modal datasets into MSMs presents both a conceptual strength and a technical challenge. For example, parameterizing a model involving concentrations of (anti-)inflammatory chemical mediators may be simple to implement if data are available (as in [17]). However, deriving credible model results at that scale without data would prove challenging. If it remains unclear whether or not additional scales are impactful, or if those scales are only of interest under specific circumstances, we apply tuneable resolution to help make a decision regarding when and whether or not to use fine-graining of additional scales (see §3.2 and [61]). Moreover, more fine-grained and computationally intensive models may struggle to yield detailed results from methods such as global sensitivity analysis (see §3.1). When adding fine-grained scales to a model, the computational demand of the smaller-scale model is increased. However, large-scale representations can be calibrated or validated by corresponding population-scale data, such as demographic data, and provide insights to large-scale dynamics such as those ranging from withinhost up to population-scale studies [53]. Lastly, we have investigated how different choices of scales and models can reveal different mechanisms [36].

In practice, to choose what information and how that information is passed between scales, we use both knowledge of a biological system and our knowledge of behaviors of model types. Linking two individual-scale models involves a continual interchange of information between individual-scale sub-models in a manner that either enmeshes their formulations in a nontrivial manner when running both models, or reduces one of them into a surrogate. For frequently-used model components, multiple frameworks including CompuCell3D [119], Simmune [81, 114], PhysiCell [40], and Virtual Cell [18, 109] have been developed to package ready-linked multi-scale models with some ability to include new components as needed. Surrogate models (i.e. simplified proxies of a model; also called emulators) must be treated with care because some may miss important fringe–cases or rare events that more fine-grained models are able to better predict, such as in the case of machine learning algorithms deployed with insufficient training data [1, 2, 3, 5, 87].

Linking time scales requires exchanging information to periodically synchronize them. For example, molecular scale diffusion, like cytokines, would have a fast diffusion rate while cellular movement or cellular updates (like death, division, etc.) happens on a much slower time scale [7, 60]. Linking these computationally can be challenging and we have reported ways that we have addressed this issue when linking information between ABMs, ODEs, and PDEs [16]. The idea in these hybrid computational models is that a computational time step is chosen based on the fastest time scales (for example molecular diffusion, for which we use 6 seconds). The simulation is synchronized when the fast time scale runs and then “catches up” where the fast time-scale meets the slower one [16, 19, 124].

With a foundation in linking scales, we can extend our discussion on formulating models to host-scale TB models where our goal is to model a single TB host experiencing multiple granulomas, as this is the typical scenario for infected primates. To do this — based on computational resources — we could not retain the fine-grained *GranSim* ABM version of granuloma formation for each granuloma, so we track each granuloma with an ODE representation. The coarse-grained granuloma scale allows us to examine a whole lung of granulomas. We created *MultiGran* for this purpose, where the ABM has as its agents individual granulomas with a system of ODEs describing each [125]. This choice allows us to address whole-host questions with fast model execution speed while affording as much biological insight as possible. We model individual granulomas with a system of ODEs describing interactions between bacteria and host cell subpopulations. Independent parameterization of those ODEs incorporated the crucial heterogeneity observed in granulomas [9, 63, 69]. The larger-scale representation of a whole-lung in *MultiGran* records a location of each granuloma, however the information passed between granulomas was limited with the exception of granuloma merging events [125]. Thus, we have a host-scale model with coarse-grained granulomas with cells, bacteria, etc. at the southern scale, and emergent host dynamics at the northern scale (Fig. 1).

In *MultiGran*, individual granulomas are endowed with independent, heterogeneous sources of non-differentiated immune T cells that allow them to grow. In a detailed study of granuloma dissemination — a process by which a granuloma can seed new granulomas — we allowed bacterial levels within a granuloma to provide a probability of dissemination [125]. To more deeply study the role of a primed immune system and trafficking of immune T cells through the blood to the lungs to participate more mechanistically, we replaced the coarse-grain representation of constant recruitment of immune T cells to the lungs by fine-graining two new physiological compartments with equations; an example of east–west scaling. This new model, *HostSim*, introduces well-mixed whole-host scale lymph-node and blood compartments [53]. These compartments are primarily driven by dynamics of ordinary differential equations that describe clonal selection and expansion of Mtb-specific CD3^+^ T cells. Unlike standalone ODE models, *HostSim’s* LN dynamics depend on antigen presenting cells (APCs) that traffic from granulomas within lungs to the LN compartment [53, 82]. Similarly, granulomas in *HostSim* no longer source their T cells from simplified recruitment functions — TNF-α and activated macrophages — but rather they recruit Mtb-specific cells directly from the blood compartment, which contains T cells effluxed from the LN compartment (Fig. 2). We also gain the ability to instantiate new granulomas as the result of dissemination events, each containing its own ODE trajectory driven by similar parameters. Our result is a host-scale model where APCs (cell-scale) sent from each granuloma (tissue scale) will affect T cell availability of other granulomas (whole-host scale). Conversely, the availability of T cells (whole-host scale) affects the ability of each granuloma (tissue-scale) to respond to infection.

We have discussed methods by which we can pass information between scales in a MSM, but we may also benefit from cross-model scale-linking. We have used *GranSim* to calibrate *HostSim* in cases where experimental data were not available [82]. In this way, we can treat detailed simulation outcomes from a detailed smaller-scale MSM as synthetic data in the same way that we calibrate a model to experimental data (see §2.3). The use of synthetic data to inform models must be performed carefully, as we might accidentally use synthetic data outside of the model context. Such misuse might introduce bias into our calibration [3], so it is important to acknowledge the appropriate context of any synthetic data.

### 2.2 Parameter Space Sampling for Model Simulation

After developing a mathematical or computational model, simulation requires choosing parameter values that are biologically relevant. Sampling values in parameter space is a necessary first step. The goal of sampling parameters in multi-scale models is to recapitulate the variation observed in biological outcomes by varying model parameters. To address the large dimensionality of parameter space, we need appropriate methods to strategically sample combinations of parameters to fulfill the particular sampling purpose. For example, local sampling methods may help overcome the uncertainty of biologically feasible parameter space such as during model calibration (described in the next subsection §2.3). On the other hand, global sampling methods do not need to run model simulations to generate subsequent parameter samples and instead generate all parameter samples simultaneously. Global sampling assumes that the parameters are independent.

We have considered several global sampling methods. There are random methods such as Monte Carlo and Latin hypercube sampling (LHS) [80] and deterministic methods such as Sobol’ sequences [117]. We compared these sampling methods and found that LHS and Sobol’ sequences cover parameter space more uniformly and converge faster than random sampling [98], and we use LHS in our studies. LHS creates *N* distinct parameter sets by dividing each parameter distribution into N equal probability intervals and sampling from these intervals without replacement [73]. The choice of *N* depends on the number of varied parameters *k*; *N* should be at least *k* + 1, however, we usually choose a much higher *N* (≈ 5000) to ensure that the whole parameter space is covered.

Sequential Monte Carlo (SMC) is a popular method of local sampling for systems biology model calibration because it is one of very few methods where increasing the number of Markov Chains explores more parameter space and therefore the sampling method scales with larger numbers of model parameters [22]. We developed a calibration approach that chooses samples to better match datasets as a way to revise parameter ranges.

### 2.3 Calibrating to Datasets using our CaliPro Method

Multi-scale models often need to include parameters that are challenging to measure from biological experiments. Additionally, sometimes parameters represent a composition of mechanisms such that no single biological measurement can be used to estimate its value. An initial guess of biologically plausible values for such unknown parameters may span magnitudes, and consequently parameter ranges need to be reduced by comparing model outcomes to experimental datasets. This process of fitting parameter ranges is called *calibration*. Calibration differs from regression because it both sets ranges of unknown parameters to closely match experimental observations including their variation, rather than solely single parameter values [52]. A further challenge for calibration methods is that we perform parameter range adjustments on many such unknown parameters with associated large degrees of freedom. We outline two methods for calibration, Calibration Protocol (CaliPro) with LHS global sampling [52], and approximate Bayesian computing with sequential Monte Carlo local sampling (ABC-SMC) [93, 116], neither of which requires a likelihood function and both propose new parameter ranges using summaries of model simulations.

Typical statistical methods of fitting parameters to experimental datasets rely on the likelihood function [101], where the likelihood function is defined as the joint probability of the observed experimental datasets and the parameterized model. However, calculating likelihood is intractable for sufficiently complex models, which is the case for most multi-scale models. We work around this limitation using pseudolikelihood, which approximates the likelihood using a distance function or kernel density estimator that compares model simulation outputs to experimental datasets as shown in Fig. 3.

**Fig. 3.**
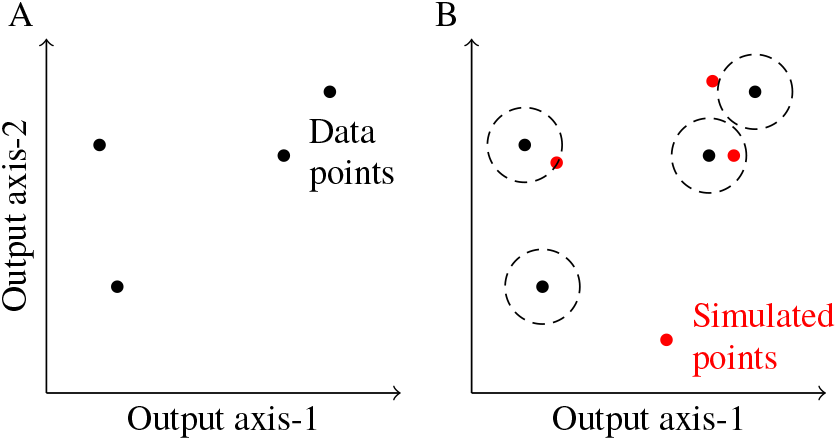
Pseudo-likelihood distance function. (A) Traditionally, likelihood is the joint probability of the data and the model parameters (B) but when calculating likelihood is infeasible, pseudo-likelihood instead considers the distance from the data to simulation output (dotted circles), where lower distance represents higher pseudo-likelihood.

We used two pseudo-likelihood based methods of multi-scale model calibration. The CaliPro approach Boolean classifies parameters of model simulations as passing or failing when simulation outputs are within or beyond relaxed boundaries of the data, respectively [52]. The passing and failing parameters are aggregated to adjust parameter ranges as well as measure the goodness of fit using a percentage scale [52] (Fig. 4A). The non-Boolean approach is to establish summary statistics to compute the pseudo-likelihood value and locally sample guided by pseudo-likelihood to set parameter distributions.

**Fig. 4.**
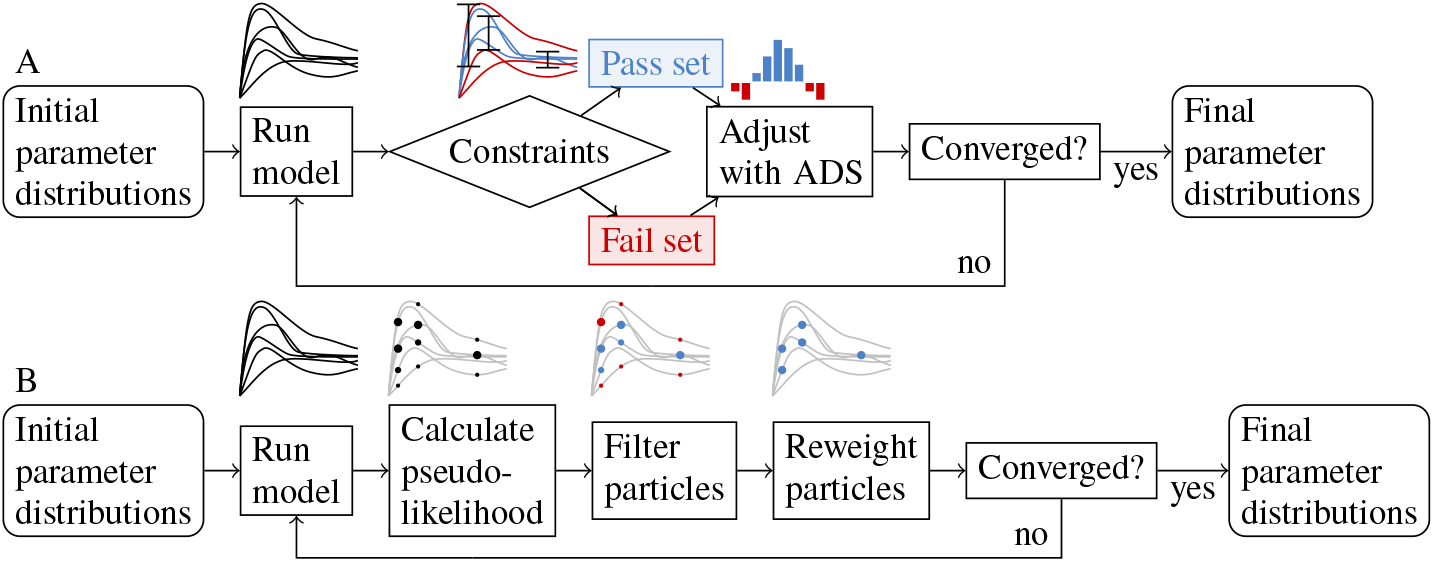
Iterating parameter ranges using Boolean and pseudo-likelihood distance summaries. (A) CaliPro uses Boolean constraints to filter parameter sets that produce higher passing rates. ABC-SMC weighs parameter particles by pseudo-likelihood to exploring parameters space with higher pseudo-likelihood. The principle of SMC sampling is assigning weights to parameter sets, filtering to parameters with higher weight, and reweighting the accepted parameter set for the subsequent iteration. Strictly speaking, particle weighting and filtering occurs in parameter space rather than outcome space, but particles have been illustrated in outcome space to aid broad understanding.

The last remaining component of these pseudo-likelihood-based calibration methods is choosing a new set of parameter ranges at the end of each iteration. Using the former approach of Boolean classification, new parameter ranges are calculated from batched global samples using alternative density subtraction, which enriches for regions of sampled parameter values that exceed the previously sampled parameter values. Collectively, this approach is called the Calibration Protocol (CaliPro) [52]. Using the approach of computing pseudo-likelihoods for each parameter set or *particle*, the pseudo-likelihood values are applied as weights to locally sampled particles and then the locally sampled particles are filtered for higher weights [67, 116] (Fig. 4B). At the end of an iterative process, the credible intervals of all accepted locally sampled particles constitute the new parameter ranges. This approach is known as approximate Bayesian computing with sequential Monte Carlo sampling (ABCSMC) [93, 116]. These concepts of comparing model simulations to experimental datasets and choosing new values is illustrated in Fig. 4.

After presenting the rationale and operation of these two calibration methods, we can compare their unique strengths and limitations. CaliPro works well with large sets of parameters by assuming parameter-independent global sampling. Formulating and composing pass-fail constraints is simpler, supports fitting a wide range of simulation outputs, and has a more meaningful and intuitive pass-percentage cut-off for deciding when to end calibration [52, 86]. In comparison, the local sampling of ABC-SMC favors uncovering multi-modal parameter distributions and parameter sets that are dependent, but has a harder time working with intractable regions of parameter space and requires formulating summary statistics [116].

## 3 Analyzing Multiscale Models

We use *analyzing* to refer to the introspective examination of models — understanding driving mechanisms and parameters, longevity of models, etc., which we hold separate from *applying* models — deriving insights into biological systems (see §4). Here, we review three model analysis paradigms to sustain credible models.

### 3.1 Uncertainty and Sensitivity

Multi-scale models involve many parameters, variable initial conditions, and stochastic events; therefore, ascribing the sensitivity of model outputs to each of these drivers is challenging. Moreover, the sensitivity to each driver varies over time in dynamical models. For model outputs of interest, assessing output uncertainties and driver sensitivities are an invaluable tool for interrogating biological questions.

Output uncertainty has two forms: epistemic and aleatory. Output uncertainty that arises from not knowing the ground truth of parameters or variable initial conditions is called *epistemic uncertainty* [47, 73]. To overcome epistemic uncertainty, we conduct sensitivity analysis (SA) to thoroughly explore outputs of the model by considering the whole biologically-relevant parameter space and variable initial conditions. We perform this either with a local sensitivity analysis, where we vary one parameter at a time, or a global sensitivity analysis, where we explore the whole parameter space simultaneously [73, 97]. Local sensitivity analysis is not as informative when applied to complex biological models, as the sensitivity of varied parameters may depend on values of all other fixed parameters [73]; however, local sensitivity analysis is used on very large models where it is challenging to compute global sensitivity indices. Hence, for our MSMs, we prefer the global approach when analyzing output sensitivity.

The second form of output uncertainty is aleatory. Deterministic models have only epistemic uncertainty, as each simulation with the same input generates the exact same output. However, stochastic models contain an inherent randomness that leads to varying outputs even with the same input. The uncertainty that arises from stochasticity is called *aleatory uncertainty* [47, 73]. To account for aleatory uncertainty, we simulate from our model repeatedly — usually at least 5 times — using the same input (i.e., same parameter set and initial conditions of variables) but different series of stochastic events, resulting in different model outputs [73].

There are several methods to quantify model sensitivity to parameters, and the appropriate method for SA depends on the model dynamics, specifically the relationship between model inputs and outputs. If the relationship is *linear* using the Pearson correlation coefficient, partial correlation coefficient or standardized regression coefficient for SA is appropriate [73]. Alternatively, if the relationship is *nonlinear*, using a rank-transformed method such as Spearman rank correlation coefficient and partial rank correlation coefficient (PRCC) for monotonic relationships, or a method based on decomposition of model output variance such as Fourier amplitude sensitivity test (FAST) and its extended version (eFAST) for non-monotonic relationships is warranted [73]. Combining uncertainty analyses with the appropriate sensitivity analyses helps us elucidate the mechanisms and factors driving model output.

### 3.2 Tunable Resolution

We developed tunable resolution: a multi-beneficial approach to fineand coarsegraining existing multiscale models [61]. Fine-graining involves refining models that are highly multiscale; however, it may be important — for computational reasons or other reasons — to reduce, or coarse-grain, a model. Usually, we create a fine-grained sub-model to simulate in more detail a phenomenon that was previously represented by a parameter or simple expression. Conversely, the goal of coarse-graining is to engineer the reverse process, but retain important information gleaned from the finegrained behaviors to improve the newer coarse-grained representation [61]. This typically applies to models that have a detailed north–south sub-scale model or an east–west sub-model. Sensitivity analysis, described in the previous subsection §3.1, is key to identifying critical model parameters to fine-grain or conflating parameters with less impact to coarse-grain [61, 73]. Note that tuneable resolution is distinct from mathematical model reduction, which focuses on coarse-graining the mathematical representation of a model for which the goal is to preserve the mathematics at the coarser scale. In the case of tunable resolution, the goal is to preserve the biology that was gleaned from the finer-scale sub-model. Tunable resolution is also distinct from multi-resolution modeling, which focuses on coarse-graining known fine-grain system dynamics. Tuneable resolution focuses on dynamically coarse-graining or fine-graining the biological representation of a system, and feeding new insights gained by one representation to the others. This process can even be automated, as we have done previously [43].

As an example of a multiscale model that we added a fine-grain sub-model, in previous work we identified that the production of the molecule Tumor Necrosis Factor (TNF) is important to the inflammation and control process in TB and is involved with long-term host outcomes [27], Given the complexity of TNF levels in granulomas, we developed a fine-grained sub-model to represent TNF production in *GranSim* [27]. Using the fine-grained model, we determined the elements of the TNF pathway that were most impactful to *GranSim*, and used the analysis to improve our coarse-grained model [61]. Consequently, we can now toggle between the two sub-model versions as needed, depending on the biological question. Later, we further increased the detail of the TNF pathway to include the transcription factor NF-κB [28, 29, 61]. This enables us to use *GranSim* to study the impact of this transcription factor, but also gives *GranSim* the flexibility of pursuing other TB-related systems of interest without spending unnecessary computational resources.

While this method is applicable to any multi-scale model, the challenges of linking scales outlined earlier still apply. For example, if our tuneable sub-models vary in their time or spatial resolution, we need to pay special attention to the mode by which our alternate sub-models couple with the rest of the model system [61]. Similarly, we must make sure our sub-models’ coupling to the rest of the model is flexible if they use different modes of computation [61]. For instance, a fine-grained sub-model could represent bulk cell motion with an agent-based model and a coarse-grained model could use partial differential equations. Therefore, paying close attention at the beginning of a project to efficiently tuning resolutions is important because it impacts the choice of modeling modes of computation and methods.

Mathematical models need to balance representing realities of biological systems against computational burden. Specifically, we need to represent biological realities as closely as possible without unduly increasing computational burden of model simulations. In spatial, agent-based models like *GranSim*, one source of computational burden is the grid dimension: 2-dimensional (2-D) models are computationally more efficient, but may lack spatial information that 3-D models provide. To investigate this issue, our group developed a 3-D version of *GranSim* to compare and contrast against the 2-D version [74]. Our study found evidence that the same mechanisms drive bacterial populations in both models. Yet, the 2-D model is more sensitive to cell density and crowding than the 3-D model, as expected, because cell movement is more limited in 2-D. Moreover, extending *GranSim* from 2-D to 3-D increased the computational burden for simulations by 2 orders of magnitude [74]. Therefore, a 2-D model is more efficient to use, unless a research question relates to cell density or cell movement. Another concern arising from the 2-D model is that the *in vivo* or clinical datasets that we use for model calibration are obtained from 3-D biological granulomas. To calibrate our parameters in a more realistic way, our group developed a methodology for scaling 2-D model outputs into 3-D to better represent biological datasets [100].

Effectively implementing tuneable resolution requires implementing models in a modular way, meaning that there are clear boundaries between model components [61]. Modular model implementation has the added benefit of more clearly delineating between various biological components represented in the model, which aids horizontally linking models [53] and clear, credible calibration of model components.

### 3.3 Ten Rules for Model Credibility

We have discussed the methods by which we develop multi-scale models. A consistent and important challenge is establishing the credibility of the model both for ourselves as modelers, as well as accuracy for our scientific and clinical collaborators. With many moving parts and potential pitfalls in the modeling process, we use ten simple rules for model credibility that were generated as part of the Multi-scale Modeling Consortium [33]. These rules, that were listed in the introduction, are generally applicable to modeling most biological and biomedical processes and they address many potential points of failure in the process of model development.The ten rules for model credibility are as follows: 1: Define context clearly; 2: Use appropriate data; 3: Evaluate within context; 4: List limitations explicitly; 5: Use version control; 6: Document adequately; 7: Disseminate broadly; 8: Get independent reviews; 9: Test competing implementations; 10: Conform to standards. There is room for broad interpretation for how these rules are implemented and may be model context dependent. Throughout our paper we highlight how we address aspects of these rules; Table 1 lists specifically how we have addressed these ten simple rules.

**Table 1.** Our implementation of the 10 rules of model credibility.

| # | Rule | Our implementation |
| --- | --- | --- |
| 1 | Define context clearly | For the models we created, we focus on the context of Mtb infection and work directly with our experimental collaborators to ensure our models are grounded in the biology specific to the questions we are studying. |
| 2 | Use appropriate data | When data are not available, we are able to sometimes have our collaborators generate those data when possible. |
| 3 | Evaluate within context | In the same way that we build models in context, we evaluate them against different data to ensure we are appropriately capturing known infection dynamics and different scales of interest. |
| 4 | List limitations explicitly | When we build a model, we always include a list of assumptions that are made about both the biology and the modeling approach we are using. In the discussion, we typically outline the limitations of the study and make suggestions for the current and future steps to address those. |
| 5 | Use version control | Our lab has used a version control method known as Subversion [91], to track both short and long term changes to code; this is especially important for *GranSim* which has been continuously curated against data since 2004[110]. |
| 6 | Document adequately | We host websites for our models that provide documentation for our models that is also continuously curated with parameters and model updates as they become available (see for example http://malthus.micro.med.umich.edu/GranSim/). |
| 7 | Disseminate broadly | We publish results of studies on our lab website, biorXiv, and in well-established journals to participate in the peer review process and to broadly disseminate our models and their predictions. |
| 8 | Get independent reviews | We have worked closely with the lab of Dr. Jennifer Linderman at the University of Michigan as well as two faculty outside of our institution, Drs. Silvia Blemker and Shayne Pierce-Cotler (University of Virginia) over the past 6 years to maintain a friendly independent review process of our models, our websites and our manuscripts. These 3 scientists together with one of us (Kirschner) presented the inner workings and results of this important collaboration[6]. |
| 9 | Test competing implementations | Interestingly, we have approached the rule of competing implementations in a number of ways. First, we have implemented in a number of ways the idea of modeling a granuloma and then used sensitivity analysis to help us identify what we learned from each approach[35]. This helped us to identify which formulation would work best for the biological questions we are posing. Next, two individuals were able to reproduce developing *GranSim* by the rules that we publish in papers and on our *GranSim* website; these groups used their versions of *GranSim* to explore different aspects of granulomas such as oxygen transport and availability within granulomas[20, 44, 45, 79]. |
| 10 | Conform to standards | Finally, by implementing the above rules into our framework and calibrating and validating our models to datasets and testing negative and positive controls of our models with known biological outcomes. We believe this satisfies the final rule of conforming to standards. |

## 4 Applying Multiscale Models: Generating Novel Outputs and Insights

With a computational model that has been calibrated against available datasets, we are now able to address questions that we are interested in and that the model has been developed to consider. There are many ways to generate results from a computational model, but each should be tempered with an understanding of model limitations, namely, not over-stating our results, when considering our necessary choices of simplification [3].

We briefly mention five examples of non-mutually-exclusive approaches of generating insights from models: (i) *in silico* experiments (ii) multi-scale interventional design studies [82] (see §4.4), (iii) analyzing uncertainty and sensitivity analyses (see §3.1), (iv) optimization investigations (see §4.2), and (v) exploratory models.

(i) *in silico* experiments refer to treating our model as though it was an accurate representation of the true system itself, and we consider simulation outputs as raw synthetic data [82, 115]. This allows us to generate a virtual cohort of sorts, by sampling our parameter space to make many virtual individuals. While this method is straightforward, it requires cautious examination of the strength of our findings. (ii) Multi-scale interventional design leverages our ability to use the same virtual subject multiple times under various conditions to determine the importance of subject heterogeneity on intervention response and vice versa [82]. Uncertainty and sensitivity analysis are powerful tools (discussed in §3.1) that allow us to determine principal model components that drive model outcomes of interest. (iv) Optimization allows us to query a system for specific maximal or minimal cases, such as ideal drug dosing or vaccination strategies based on our knowledge of a system [8, 13]. (v) Exploratory models determine which hypothesized mechanisms best explain or recapitulate data in a complex biological system [8, 13]. Below we highlight specific examples of these five approaches and how they operate for our systematic study of TB.

### 4.1 Granuloma Dissemination

We explained how granulomas are structures that form within lungs that physically contain and immunologically restrain bacterial growth (see §2). Primate hosts have a median of 14.5 granulomas per infection with Mtb [8, 125]. In at least half of hosts, some granulomas completely clear their bacterial load, including hosts that suffer active TB disease [65]. In some cases, granulomas that do not clear their bacterial load can disseminate and seed a new granuloma. This can happen either locally — next to an existing granuloma, which we refer to as *offspring* granulomas — or distally — even as far as to the other lung lobe. Mechanisms that lead to granuloma dissemination are unknown and include events such as neutrophil seeding, structural breakdown due to bacterial overgrowth, spilling of bacteria into airways, etc.

In §2.1, we reviewed our whole-host model for TB, *HostSim*. In our first introduction to the *HostSim* model, we represented granuloma dissemination events as follows. At every time step in the model we assumed that there is a chance for any granuloma to disseminate and that the dissemination probability increases as that granuloma’s Mtb levels (colony forming unit; CFU) count increases. Recent data from NHPs suggest that granulomas detected before four weeks post infection are primary granulomas [41]. Granulomas thought to be a result of disseminating primary granulomas were detected only after four weeks post infection. Shown in Fig. 5A-B are data obtained from [41]; these later-detected NHP granulomas have lower CFU counts than the earlier-detected set at the time of necropsy. We capture this trend in *HostSim* by comparing simulated disseminating granulomas to primary granulomas at 70 days post infection, allowing comparison to the dataset from ten week NHPs (Fig. 5A-D).

**Fig. 5.**
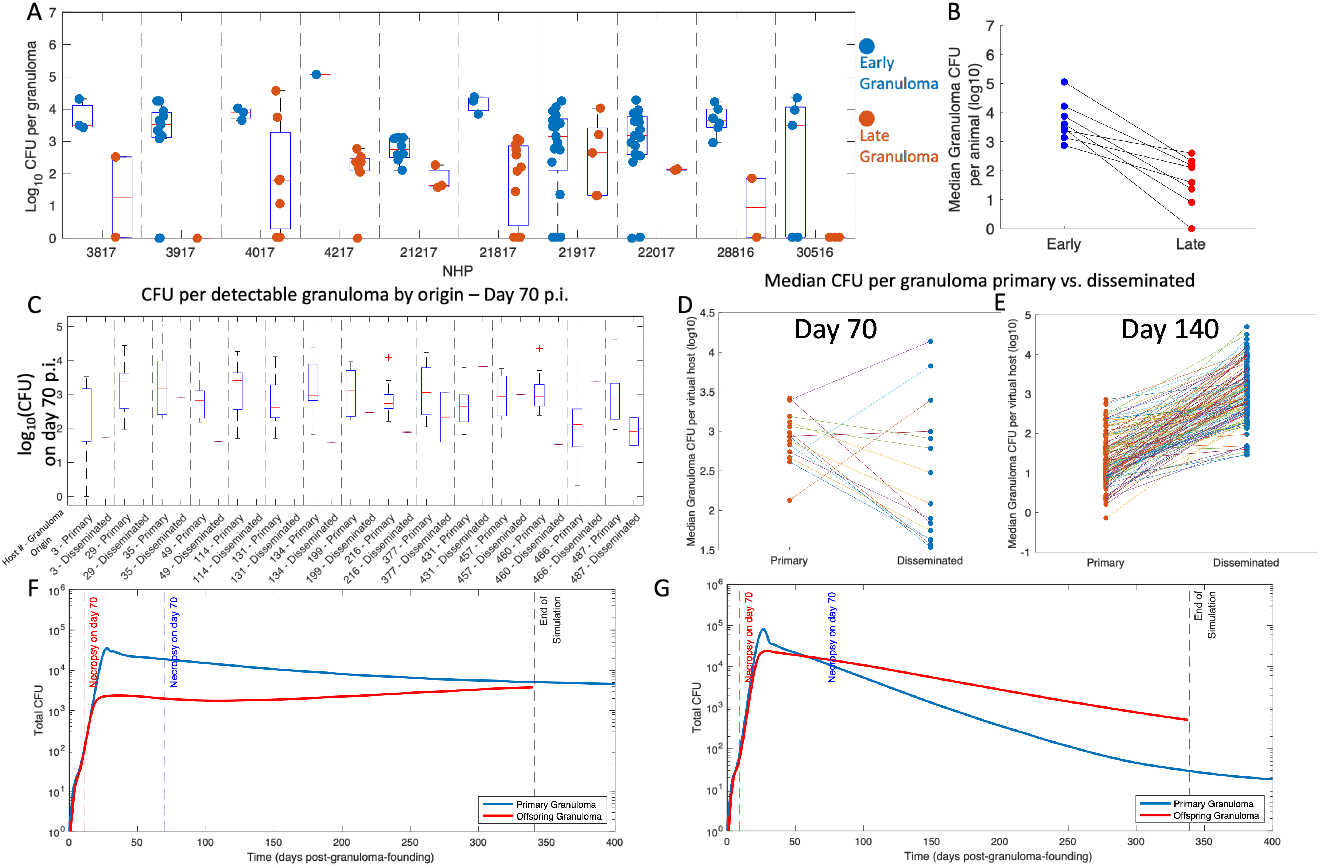
Exploring granuloma dissemination between experimental and synthetic granulomas. (A-B) Datasets from [41]. (A) Box plots of log_10_ (CFU) in early (blue) and late (red) detected NHP granulomas with the NHP numeric identifier under each set of box plots. (B) Median CFU of each animal is lower in late detected granulomas. (C) Pairs of box plots showing summary statistics of log_10_ (CFU) for (left) primary and (right) for disseminating granulomas. We obtained these data from virtual hosts on day 70. (D-E) Direct comparison between medians of primary and disseminated CFU per host on days (D) 70 and (E) 140 post-infection. (F) CFU in time post-granuloma formation is shown for (red) a disseminated granuloma and (blue) the primary granuloma that disseminated it. For convenience, we show the time of virtual necropsy (70 days post virtual-infection) as a blue dashed line, and the corresponding time of virtual necropsy as a red dashed line.

While it becomes intractable to study progressively worse granuloma dissemination in NHPs, we are able to use *HostSim* to simulate disseminated granulomas and predict their later-time dynamics at times that continue beyond the post-necropsy times of corresponding NHP datasets. Interestingly, *HostSim* predicts that while at the earlier 70-day time point the newly disseminated granulomas have lower CFUs, at later time points many of those disseminated granulomas become markedly higher-burden than the median primary granuloma within a host (¿ 140 days post-infection) (Fig. 5E). When we compare disseminated granulomas to their parent granulomas, we find, as expected, that their initial growth is much slower — consistent with the presence of effector T cells in the lymph node (Fig. 5F). However, in some cases, over long time periods, primary granulomas are surpassed by their own offspring by CFU as a function of time-post-granuloma-founding (Fig. 5G). In fact, disseminated granulomas in *HostSim* tend to be worse in the long run; performing the same analysis on the same population at 140 days post-infection yields the opposite trend. By 140-days post infection, median CFU is higher in disseminated granulomas than in primary for most hosts (Fig. 5E). This may follow because in *HostSim*, locally disseminated granulomas tend towards having parameters (behaviors) similar to their associated primary granulomas, which are themselves sufficiently high-burden to disseminate. This is even more likely because we find that even though the median CFU burden of the disseminated granulomas were higher than the median CFU burden of the primary granulomas, only 3% of disseminated granulomas were higher burden than their own parents by 140 days post-infection. This number increased to 14% by day 400 post-infection. This example highlights how modeling can extend experimental studies by performing simulations that are not currently possible (providing an exploratory result); in this case after an animal is necropsied we can simulate infection out for a longer time. We predict if that were the case, although it appears that those disseminated offspring granulomas did better at controlling bacteria, the model shows that was a short-lived phenomenon that could not have been detected experimentally. This type of *in silico* experiment is an invaluable resource for traditional wet lab experimental studies.

### 4.2 Drug Regimen Pharmacokinetics and Pharmacodynamics

Although testing interventions using conventional clinical trials is essential and informative, it is also a tedious process that takes many years and costs millions of dollars [111]. In addition, many complex diseases require combination therapy [57, 84], increasing the number of possible interventions to a level where testing each possible combination clinically or experimentally is not feasible [13]. We can use MSMs to efficiently screen multiple possible interventions [8, 14]. As pointed out above (see §4), multiscale intervention design studies are a key feature application of MSMs.

One of the most common interventions for infectious disease studies is the administration of drugs. To understand mechanisms of drug dynamics, we implement drug pharmacokinetics (PK) and pharmacodynamics (PD). PK investigates what host pathology does with drugs, namely, how a drug is absorbed, distributed, metabolized, and excreted within the human body. PD investigates what drugs do within a host, namely, therapeutic effects of drugs depend on drug concentration at sites of action. The interplay between PK and PD determines the outcome of the therapy.

Although modeling PK typically involves using compartmental ODEs that represent different physiological compartments of the host or tissue [39, 50], in some conditions, the heterogeneity of drug distributions may be critical to predict treatment outcome. For example, the complex and heterogeneous structure of TB granulomas (or tumors as well) prevent effective diffusion of drugs into the necrotic core, where bacteria are trapped [106]. Drug diffusion into the granuloma necrotic core (caseum) is a key challenge of TB treatment. Therefore, we need to consider spatial aspects of granulomas while modeling PK of TB drugs. *GranSim* simulates the formation of granulomas (see §2.1). Using *GranSim*, we incorporate a drug scale, namely PK/PD, to test various drug regimens and to help predict more mechanistically what drives drug outcomes. We included a plasma PK compartmental ODE model [89, 90] (Fig. 6A). Drugs within plasma then permeate into lungs, i.e., the *GranSim* environment, through vascular sources modeled in *GranSim* via a flux-based equation [89, 90] (Fig. 6B). Once in *GranSim*, drugs diffuse, bind to macromolecules, and partition intracellularly into immune cells (Fig.6C). These processes control how well drugs can penetrate granulomas [89, 90]. By combining *GranSim* with a PK model, we can determine distributions of drug concentration within granulomas to simulate drug penetration [89, 90].

**Fig. 6.**
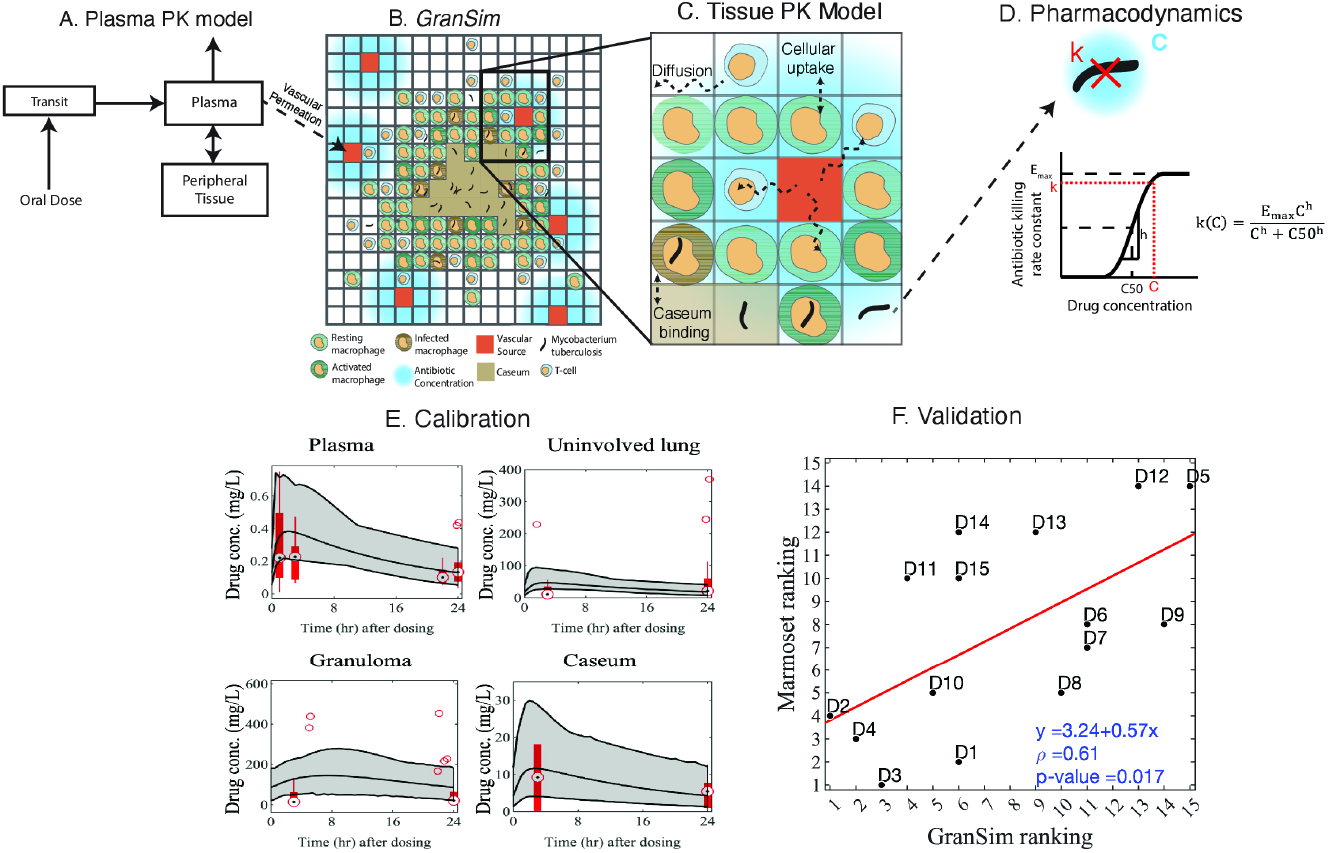
An overview of our TB treatment simulation pipeline using a pharmacokinetics (PK) / pharmacodynamics (PD) model incorporated into *GranSim*, our multi-scale model that simulates TB granuloma formation and disease progression. (A) We simulate drug concentrations in plasma using a compartmental ODE model. (B) The drug in the plasma penetrates into lungs, i.e., the *GranSim* environment, through vascular sources. (C) The penetration of the drug into the necrotic core of granulomas is determined by various tissue PK processes, such as diffusion, caseum binding, and cellular uptake. (D) We model PD using a Hill equation and determine the drug killing rate constant based on the drug concentration at every grid compartment. (E) Calibration of an example drug to *in vivo* data from various tissue samples, such as plasma, uninvolved lung, granuloma, and caseum. In each panel, red box plots represent *in vivo* data for an example drug. The dots in the middle of the box plots represent the median, whereas the top and the bottom of the thick red column of the box plot represent the 75th and 25th percentile, respectively. The black lines are the maximum, mean, and minimum drug concentrations from *GranSim* simulations, and the gray shaded area is the area between the maximum and the minimum concentrations. Only PK calibration is shown in this panel. (F) Once parameters for all drugs of interest are calibrated in *GranSim*, we then simulate all drugs, rank them, and compare these *GranSim* rankings to experimental drug rankings — animal model or clinical trials (D1–D15: 15 different drugs or drug combinations; correlation was determined using Spearman’s rank correlation).

Many PD models have been derived that predict drug effect based on drug concentration, however, the model we found that best supports our experimental data is the Hill equation [70]:

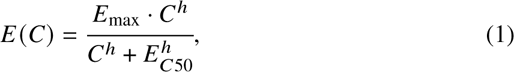

where *E*, our drug effect, is a function of drug concentration, *C*, and depends on parameters *E*_max_ (maximum drug effect) and *E*_*C*50_ (the concentration of the drug needed to achieve half maximal effect, *E*_max_/2). In *GranSim*, we determine the number of killed bacteria within each grid compartment using a Hill equation and the distribution of drug concentration is assessed by the PK model [89, 90] (Fig. 6D). Calibrating PK/PD model parameters to experimental data is a crucial process to represent each drug in a realistic way. In *GranSim*, we model PK using parameters such as how much drug penetrates into the caseum, diffusivity, vascular permeability, degradation rate constant, intracellular-to-extracellular drug concentration ratio, etc. Some of the PK parameters can be directly measured *in vitro*, such as caseum penetration (unbound fraction) [106] and intracellular-to-extracellular drug concentration ratio [104]. However, the remaining parameters are almost impossible to measure *in vivo* or *in vitro*. Thus, we calibrate these unknown parameters using *in vivo* drug concentration datasets from humans [92] or animals that develop necrotic granulomas like humans, such as non-human primates [122] or rabbits [128] through our systematic calibration process, CaliPro (see §2.3) (Fig. 6E). We then validate our calibrated model using MALDI mass spectrometry images of drug penetration within a granuloma [50]. To calibrate PD parameters (parameters of Hill equation, i.e., *E*_max_, *E*_*C*50_, ℎ), we use *in vitro* bactericidal assays that are performed with antibacterial drugs [23, 105]. We then validate our predictions of drug efficacies to clinical trials [14] or NHP efficacy studies [8] (Fig. 6F). To do that, we first rank the regimens (annotated as *D*_*i*_ in Fig. 6F) based on results from clinical trials or NHP studies (experimental drug rankings). Then, we simulate the same regimens in *GranSim* and rank them based on the simulation outputs (*GranSim* rankings). Finally, we calculate the correlation between experimental rankings and *GranSim* rankings (Fig. 6F).

Our GranSim-PK/PD pipeline (Fig. 6) has successfully predicted clinical trials and NHP experiments [8, 14] proving the credibility of our approach. However, for diseases that require a combination therapy like TB, the number of possible combinations to test may exceed computational limits and resources [13]. Therefore, we incorporated an optimization pipeline to accelerate the process of testing by making wiser predictions and we discovered new combinations that haven’t been tested clinically yet [8, 13]. MSMs combined with an optimization pipeline will expedite the discovery of new combination regimens by informing experimentalists with decision-making results.

### 4.3 Vaccination

A key aspect of building models that capture immune response to infection is that we have the opportunity to study vaccination. Vaccination is another key TB intervention that is currently offered in some, but not all, countries. This widely used Bacillus Calmette-Guérin (BCG) vaccine offers a range of 0–80% protection in both infants and adults [4, 120], and is not administered in the USA and the UK, due to this variable efficacy. Presently, a number of TB vaccine candidates are being actively developed with more than 20 entering clinical trials and 14 being actively evaluated [59]. Below, we illustrate two studies we performed in this area: the ideas we have developed around the concept of a vaccine design space [129] and how we explored mechanisms responsible for key differences between NHP and human vaccine responses in TB [54].

We explored the idea of creating a vaccine memory design space where our goal is to identify protective levels of CD4^+^ and CD8^+^ memory T cells and, more recently, tissue resident memory T cells [51, 129]. Memory design space visually separates antigen-specific memory cells from their functional properties to help identify antigen-specific and antigen-independent subpopulations that confer protection [129]. In previous modeling work, we identified how persistence of different T-cell types as a percentage of their peak population during infection related to disease outcomes of latency and sterilization [118]. Additionally, memory responses from synergistic combinations of these T-cell types require substantially fewer numbers of each type of cell, predicting the need to target both CD4^+^ and CD8^+^ cells to develop an efficient vaccine [118]. Effector memory T cells circulate through blood whereas central memory T cells circulate through lymph nodes and are longer-lived. Besides a need to target generation of both CD4^+^ and CD8^+^ cell types, our study hypothesizes that vaccine efficacy is also dependent on different levels of effector and central memory cells for each antigen-specific memory population [129]. Therefore, the vaccine design space concept predicts vaccine formulations that generate synergistic antigen-specific memory T cell populations.

Whereas the previous studies assumed tissue resident memory T cells are a subset of effector memory T cells [31, 129], we had a more recent study that explicitly models tissue resident memory cells in *HostSim* [51]. This allows us to investigate outcomes during a re-exposure event that occurs concomitantly during primary infection [51, 53]. Unlike previous vaccine studies that vary numbers of vaccine doses over a few months [54, 129], we simulated reinfection of virtual hosts in the tissue resident memory T cell study for up to 7 years [51], illustrating the usefulness of MSMs to capture long-term protection from secondary infection that would have been both slow and expensive to study with animal models.

We also further investigated a vaccine, known as H56, a recent TB vaccine candidate with potential for long-term immunity [54]. For this study we focused on immunogenicity of H56 and simplified our model using our biological knowledge of the H56 vaccine failing to induce a robust CD8^+^ T cell response [68], by using a 2-compartment model of the lymph node and blood with 16 nonlinear ODEs [54]. Our model predicted that the third dosing was not necessary as it did not provide any additional protection. Had results from a model like ours been available prior to the clinical study, the information would have saved time and resources of researchers and patients. Additionally, by calibrating the same model separately against both NHP blood data and human blood data [68], our model sensitivity analysis was able to validate mechanistic appropriateness of NHPs as a model following *in vivo* comparisons [32, 55, 88, 108]. One key difference with the human system was observed— that antigen presentation to T cells in humans was not as effective as compared to antigen presentation in NHPs [54]. This mechanistic insight was uncovered from significant negative correlations of half-saturation values of proliferation and differentiation, where the half-saturation values represent the likelihood of proliferation or differentiation after T cell priming [54]. This highlights the ability of sensitivity analysis to isolate mechanisms leading to distinct vaccine responses and to evaluate accuracy of using model systems to capture human disease.

### 4.4 Multiscale Intervention Design for Disease

Once we have adequately represented an intervention (e.g. a drug treatment or a vaccine), the next question is how well an intervention works. Classical analyses often investigate treatment group summaries of patient cohorts. While this captures the efficacy of an intervention at the scale of a group of patients, aggregating patient data mixes effects of patient heterogeneity and also may cloud details occurring at within-host scales. To avoid losing patient-specific intervention effects, we developed a multi-scale interventional design framework that compares virtual patients against themselves in both control and intervention scenarios [82]. This method generates a cohort of virtual patients that can be represented by both control and intervention scenarios (models), and a biologically transparent method for comparing outcomes between those scenarios on a sub-patient scale that we refer to as an intervention efficacy [82].

TB has a high level of intra-patient heterogeneity [9, 63, 69] and this will benefit from applying a multi-scale interventional design framework [82]. With this framework, we can digitally represent a collection of patients both with and without treatment and have an *in silico* ideal negative-treatment control on a per-patient basis. Collecting and quantifying effects for each patient in the intervention group will allow us to directly study the spectrum of patient outcomes, and identify (i) which patients respond best to treatment, (ii) which mechanisms are responsible for those differences, and (iii) what modes of data may be the best predictors of patient-response to intervention.

## 5 Future Directions

Multi-scale modeling is a powerful tool for understanding complex disease dynamics over multiple physiological scales that amplifies experimental datasets to generate novel insights specific to a given disease. We have developed approaches like tunable resolution and tools like CaliPro to build and improve MSMs and have used our models of *M. tuberculosis* infection to perform virtual experiments that bypass technical limitations of physical experiments and generate new testable hypotheses ranging from fundamental immune cell biology to disease-specific drug and vaccine dosing. When partnered with appropriate experimental approaches and datasets, MSMs allow us to investigate questions inaccessible to traditional approaches.

Our group has had success developing infection progression models and feeding back our insights into new studies through our long-standing collaborations with experimentalists. These collaborations allow us to not only have a greater say in the type of data collected but also help us pay close attention to feedback to ensure our models are properly rooted in our evolving understanding of the underlying known biological mechanisms. While these collaborations have been indispensable to our work, in general, researchers need more incentives and opportunities to take the time to foster these connections. We can achieve this by increasing funding for collaborations between experimental and computational researchers and institutionalizing the knowledge and practices that expand these collaborations.

Newer, more detailed experimental and synthetic datasets drive the need to revise existing models to incorporate them. Updating models with new datasets is a time-consuming, intensive task of balancing the needs of (i) including sufficiently detailed biological representation through the addition of fine-grained model components; and (ii) managing model complexity through coarse-graining the biological system. Another time-consuming task is model re-calibration that arises from significant model revisions. Sensitivity analysis is one useful tool for performing both of these tasks by identifying model parameters driving outputs. Still, we see many opportunities to expedite the process of model updates.

In addition to forming collaborations and using datasets that emerge from them, sufficiently cross-training researchers in biological, mathematical, and computational fields will seed future breakthroughs. There is a knowledge gap between mathematics or computational researchers and their biological counterparts that requires cross-training at all career levels [83].

Composing more complex MSMs would benefit from sharing and linking MSMs across platforms and programming languages. In theory, this would be optimally accomplished by a standardized, universal platform, which allows easy linking of sub-models, both vertically and horizontally. Platforms such as BioSimulations [112] aim to recreate published models using a common format to enable vertical and horizontal connections of various models. Modeling publication processes and practices can encourage greater adoption of sites like BioSimulations for compatible model frameworks, and reduce the need for models written with in-house codes that are not easily composable.

Optimization processes applied to MSMs will produce impactful interventions. MSMs have been informative for modeling interventions and predicting their outcomes [8, 14, 89]. However, for diseases that require combination therapies, the number of possible combinations to test may exceed computational limits and resources [13]. Therefore, machine learning-based optimization algorithms may be useful to accelerate the process of testing and optimizing interventions using MSMs by making robust predictions [8, 13]. If supported, the collaboration of experimentalists, modelers, and data scientists will significantly expedite clinical trials for new and more optimal interventions of diseases requiring combination therapy.

The ability to create biorepositories to store collections of virtual hosts along with their responses to various interventions will become a valuable resource in the future. Similar to digital twins, we will be able to use these repository cohorts to produce, for a specific clinical patient, a family of digital partners — those virtual patients that are measurably closest to the real patient. This will provide a valuable predictive tool — a forecast cone for each potential treatment available to a patient, along with a mechanistic explanation for that particular patient’s response. This family of close digital partners can narrow as more types of data are fed into the system, closing the gap between the partners and the patient. At the very least, it will highlight biases in the methods by which we quantify the efficacy of impact.

Another way that virtual hosts can potentially be used is to perform virtual preclinical trials. By definition, clinical trials aim to assess whether a given intervention (the factual) is an improvement over what would have happened if the intervention did not exist (the counterfactual) [25]. Given that we are unable to take a patient back in time, we assume that we can construct two groups that, when looked at as a whole, represent the same average person [25]. The major pitfall of this approach is that it assumes that patient groups can only be matched on known traits relevant to outcomes of interest. Virtual hosts provide a unique opportunity to have a perfect counterfactual as we can, in essence, take our patients back in time and apply different interventions for comparison [82]. Further, exploring treatments in this virtual sphere does away with the same ethical or loss-to-follow-up concerns that plagues traditional clinical trials [103, 107] and also would greatly reduce cost and time necessary by multiple orders of magnitude.

When time is of the essence, as in the case of rapid deployment of complex epidemiological MSMs, we need a robust pipeline for successful identification of new treatments and other means of combating infectious disease. Following the identification of a novel disease or condition of public health concern, there is an inherent need to work on multiple scales during the initial investigation into how the disease is affecting the public and during the determination of the optimal interventions to implement [56]. MSMs enable us to quickly test and examine how interventions applied at one scale impact others scales, or to identify which features of a system are important to target [96]. This information can be quickly taken to policy makers and used to inform public health interventions as was the case during COVID-19 where models were rapidly deployed and used to inform the establishment of quarantining and other interventions [96, 121].

## Acknowledgements

This research was supported by NIH grant R01 AI50684 (DEK) and is supported in part by funding from the Wellcome Leap Delta Tissue Program (DEK). Fig. 2 was created using BioRender.com.

